# Hybrid sequence-structure based HMM models leverage the identification of homologous proteins: the example of class II fusion proteins

**DOI:** 10.1101/379800

**Authors:** R. Tetley, P. Guardado-Calvo, J. Fedry, F. Rey, F. Cazals

**Affiliations:** Inria, Université Côte d’Azur, France; Unité de virologie structurale - Institut Pasteur Paris, and CNRS UMR 3569, France; Unité de virologie structurale - Institut Pasteur Paris, and CNRS UMR 3569, 25-28 Rue du Docteur Roux, F-75724 Paris Cedex 15, France; Inria, Université Cote d’Azur. 2004 route des Lucioles, F-06902 Sophia Antipolis

## Abstract

We present a sequence-structure based method characterizing a set of functionally related proteins exhibiting low sequence identity and loose structural conservation. Given a (small) set of structures, our method consists of three main steps. First, pairwise structural alignments are combined with multi-scale geometric analysis to produce structural motifs i.e. regions structurally more conserved than the whole structures. Second, the sub-sequences of the motifs are used to build profile hidden Markov models (HMM) biased towards the structurally conserved regions. Third, these HMM are used to retrieve from `UniProtKB` proteins harboring signatures compatible with the function studied, in a bootstrap fashion.

We apply these hybrid HMM to investigate two questions related to class II fusion proteins, an especially challenging class since known structures exhibit low sequence identity (less than 15%) and loose structural similarity (of the order of 15Å in lRMSD). In a first step, we compare the performances of our hybrid HMM against those of sequence based HMM. Using various learning sets, we show that both classes of HMM retrieve unique species. The number of unique species reported by both classes of methods are comparable, stressing the novelty brought by our hybrid models. In a second step, we use our models to identify 17 plausible HAP2-GSC1 candidate sequences in 10 different drosophila melanogaster species. These models are not identified by the PFÅM family HAP2-GCS1 (PF10699), stressing the ability of our structural motifs to capture signals more subtle than whole Pfam domains.

In a more general setting, our method should be of interest for all cases functional families with low sequence identity and loose structural conservation.

Our software tools are available from the `FunChaT` package of the Structural Bioinformatics Library (http://sbl.inria.fr).

## 1 Introduction

### Function prediction from sequence and/or structure

The structure - function paradigm stipulates that it is the structure (and the dynamics) of proteins which accounts for their function. The prediction of function from sequence and/or structure data is therefore of paramount importance. The search of sequence and/or structural homology may be tackled at three levels, namely for whole proteins, protein domains, and protein motifs within domains. Indeed, different functional constraints typically apply to different regions of the proteins, and even more, within a domain, internal and surface regions undergo different selection pressure depending on their involvement in the structure and/or function. In searching for such features, a relatively simple case is that of proteins harboring high sequence identity, say above 30%[1]. In that case, pairwise sequence alignments are generally sufficient to infer homology. The situation deteriorates below the previous threshold, as pairwise sequence alignments fail to capture evolutionary relationships known on the base of structure and function, especially for sequence identity less than 20%[2]. The identification of distant evolutionary relationship is generally referred to as remote homology detection, a problem usually tackled using three classes of methods, namely alignment methods, discriminative methods, and ranking methods [3]. Alignment methods resort to multiple sequence alignments [2], position specific profiles [4] as well as profile hidden Markov models (HMM) [5, 6]. Discriminative methods treat remote homology detection as a supervised classification problem aiming at training a classifier-see the numerous references in [3]. Using classical protein classifications such as `SCOP`, proteins in the same superfamily but not in the same family are considered remote homologous proteins. Finally, ranking methods approach remote homology detection as a database search. Upon embedding known protein structures into a (generally fixed dimensional) space, the query protein is used to retrieve the nearest neighbors-see again [3] and the references therein.

Because protein structures and functions tend to be more conserved than sequences [7], the use of combined sequence - structural information also offers various routes to improve the detection of remote homologs. These methods target different features of proteins, including folds, pockets and clefts, active sites, interfaces and protein - protein interactions [8, 9, 10, 11]. These methods are useful but require care. First, no best structural alignment scheme exists, and different methods typically trade the length of the alignment against its quality [12, 13]. Second, selected protein regions-loops in particular-may not have a unique structural alignment, so that purely geometric approaches may loose important sequence information. Finally, the stringency thresholds used for sequence and structure information must be chosen with care, as restrictive thresholds may generate information loss on homology distant molecules, while loose ones may yield a signal dilution [8]. Despite these features, combined sequence - structure based methods proved instrumental in enhancing remote homolog detection. Of particular interest are hybrid multidimensional alignment profiles (HMAP, [8]), a method combining primary, secondary and tertiary structure information. Under suitable hypothesis, HMAPs enhanced SCOP superfamily and fold detection [8], and it was shown that sequence and structure contain some unique information.

### Structural homology for viral fusion proteins

As an illustration of the difficulties faced when performing sequence-structure studies and remote homology detection, we consider (class II) fusion proteins [14, 15, 16, 17, 18, 19]. Membrane fusion is key to fundamental mechanisms required across the kingdom of life, from viruses to eukaryotes. For the former, viral infection requires fusing their membrane with that of the target cell. For the latter, various mechanisms involving both the plasma membrane and also internal membrane require fusion events [20]. We note in passing that commonalities of membrane fusion proteins between viruses and eukaryotes is also especially interesting: on the one hand, the functional convergence is such that structural homology is expected; on the other hand, the evolutionary pressure faced by viruses and their much shorter life cycle calls for the implementation of escape strategies that may expand the palette of mechanisms used.

In any case, catalyzing the fusion of two lipid bilayers is a complex process tightly regulated, requiring in particular the desolvation of the first hydration shell of the layers to be fused. This step is usually followed by the formation a hemifusion stalk, and finally by the pore formation. Despite their variety (class I, II, III), it is believed that fusion proteins follow the same generic mechanism. Evidence for this belief relies on the spatial proximity of the fusion loop targeting the cell membrane and the transmembrane anchor attaching the protein to the virus envelope. While this proximity in all known postfusion structures strongly hints at a common mechanism, it does not delineate the mechanism.

To further our understanding, a natural route consists of identifying commonalities between these proteins. Alas, structural studies on postfusion structures are especially challenging for two reasons [21]. First, the sequence identity is very low (less than 15%). Second, global structural alignments yield mild lRMSD (or the order of 15Å), blotting out smaller and more conserved regions. These statistics caliber the difficulty of the endeavor aiming at unveiling conservation thresholds, both for the sequence and the structures, corresponding to structural features accounting for the biological function i.e. fusion.

Fusion proteins therefore constitute a perfect case study to jointly track sequence and structure similarities.

### Contribution

We present a method performing a functional characterization of a set of proteins with low sequence identity and loose structural conservation, which delivers profile HMM biased by structural features. Similarly to HMAPs [8], our method combines structure and sequence information; yet, it bears major differences. First, the method relies on structurally conserved motifs that may span SSE and loops, as opposed to secondary and tertiary structure elements. Second, it involves a unique parameter, used to tune the stringency threshold of structural information termed relevant - see above our discussion on stringency thresholds. Third, our structural motifs are used to produce profile HMM biased towards structurally conserved regions. These HMM are then utilized to query databases such as `UniProtKB`, in order to retrieve the sequences of proteins which may exhibit the function of interest.

We validate our hybrid profile HMM on two problems related to class II fusion proteins. First, using variable learning sets involving viruses and eukaryotes, we show that our hybrid HMM retrieve from `UniProtKB` species which are beyond reach for models based on sequences only. Second, using three HAP2 structures, we show that our models identify from `UniProtKB` remote homologs in drosophilia malanogaster, which are not retrieved using the relevant PFAM family.

## 2 Material

### Viruses

As organisms which only replicate inside the living cells of other organisms, enveloped viruses use a lipid bi-layer (the envelope) to protect their genomes during extra-cellular transport to a new host cell. Initiating infection requires fusing the envelope to the host cell membrane resulting in delivering the viral genome inside the host cell cytoplasm. It is therefore straightforward why understanding the fusion mechanism is such an important stake. This mechanism is mediated by fusion proteins, grouped in three classes (I, II and III). Class II fusion proteins, which are scrutinized in this paper, are elongated molecules with three domains (DI, DII, DIII) composed primarily of *β*-sheets. The central DI domain connects via a flexible hinge to the longer DII. Typically, DII contains several conserved disulfide bonds as well as the so-called fusion loop at its tip. Additionally, a linker region connects DI to the DIII domain, which has an Immunoglobulin (Ig)-like fold. From its pre-fusion monomeric conformation, the class II fusion protein arranges itself in a trimer (post-fusion conformation) which spikes the host cell with its fusion loop. In this study, we use 6 viral fusion proteins that range from four different families of viruses: Togaviridae, Hantaviridae, Phenuiviridae and Flaviviridae. Hantaviridae and Phenuiviridae belong to the same order Bunyavirales. The others do not currently have an assigned order (Fig. 1).

**Figure 1:**
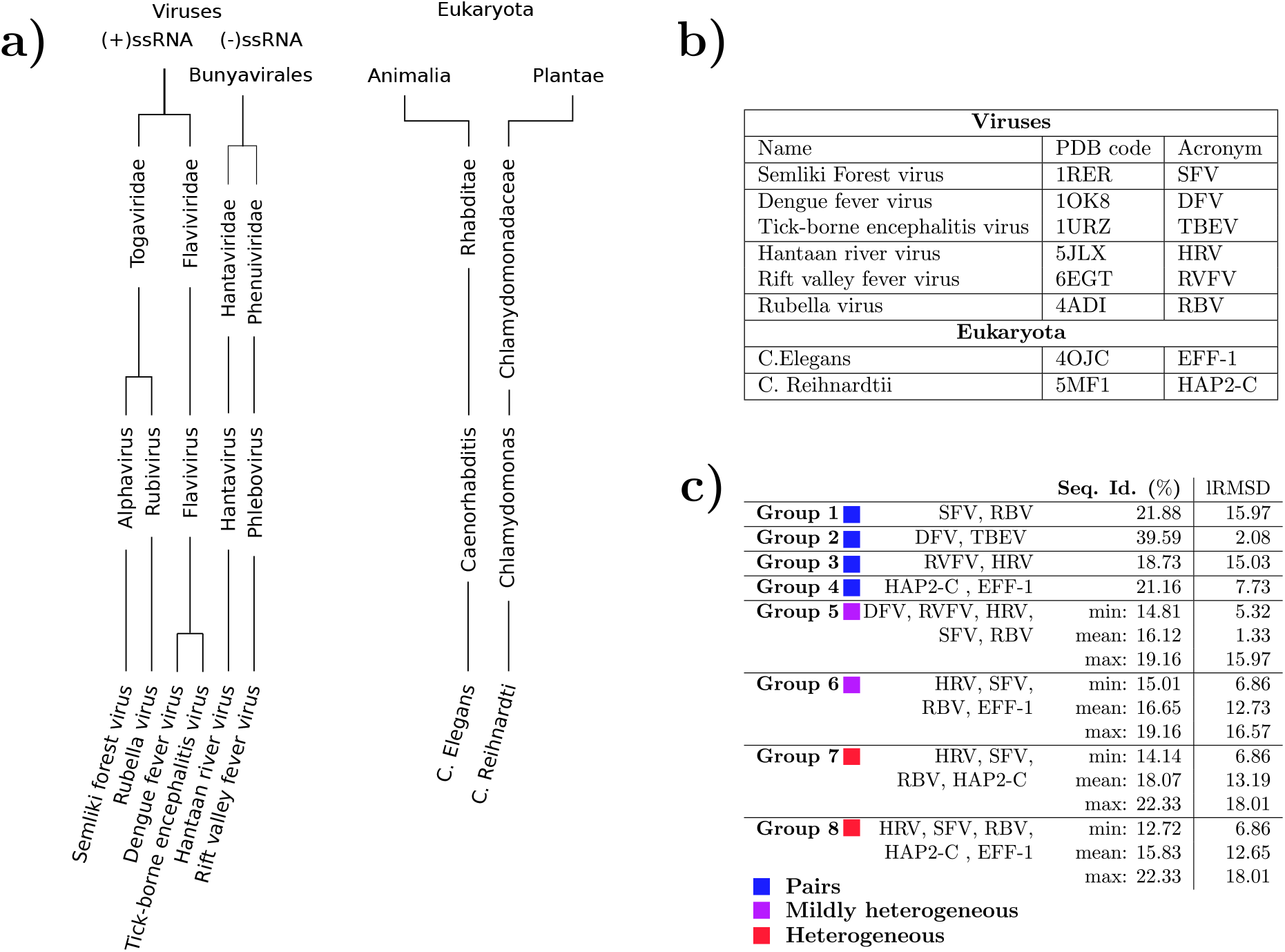
Structures used in this study. **a)** Embedding of each structure in their respective taxonomic tree (one for viruses and one for eukaryotes). We only detail the names for the genus and family ranks. The viruses are arranged in groups and the eukaryotes in kingdoms. **b)** Here we provide the files used in the study as well as the acronym used for each structure throughout this article. **c)** The groups of structures as presented in Sec. 2. For each group, we display pairwise sequence identity statistics as well as structural similarity. Regarding sequence identity, we denote three cases (which are color coded): pairs of structures (for which there is only one value), mildly heterogeneous groups (with a small interval of sequence identity values) and heterogeneous groups.

### Eukaryotes

Recent work allowed the identification of two class II fusion proteins in Eukaryotes: EFF1 [21], HAP2 [22, 23]. We add all the known crystallized structures to our study which include a nematod (C. Elegans), and an algae (C. Reihnardtii) (Fig. 1).

### Groupings

As outlined in Introduction, our method extracts conserved structural motifs so as to produce biased profile HMM. In order to assess the incidence of these conserved motifs and the resulting HMM on the the sequences retrieved from `UniProtKB`, we run our methods on various groups: **Group 1:** All Togoviridae (2 structures). **Group 2:** All Flaviviridae (2 structures). **Group 3:** All Bunyavirales (2 structures). **Group 4:** All Eukaryotes (2 structures). **Group 5:** One of each kind of virus genus wise (5 structures). **Group 6:** EFF-1 and selected viruses (4 structures). **Group 7:** HAP2-C and selected viruses (4 structures). **Group 8:** One member of each taxonomic family (5 structures).

Each of these groups is scrutinized with respect to their overall sequence identity and structural similarity. We report the minimum, mean and maximum value of sequence identity and lRMSD for each pairwise comparison in a group. We form three super-groups (Fig. 1(c)):

- **Pairs:** there are four groups (1-4) which contain only two structures. Among the four, group 2 deserves a mention for its very high sequence identity (39.59%) and very low lRMSD (2.08). Flaviviridae are very conserved.
- **Mildly heterogeneous:** Group 5 and 6 have a medium variability in sequence identity ([14.81%, 19.6%] and [15.01%, 19.16%] respectively).
- **Heterogeneous:** Group 7 and 8 are the most heterogeneous groups when it comes to sequence identity ([14.14%, 22.33%] and [12.72%, 19.16%] respectively).

More specifically, **Groups 1-5** use taxonomic groupings, while **Groups 6-8** maximize heterogeneity by balancing members of the two similarity groups.

## 3 Method: hybrid profile HMM design and database search

### 3.1 Overview

We characterize a set of functionally related proteins with known structures in three steps (Fig. 2):

**Figure 2:**
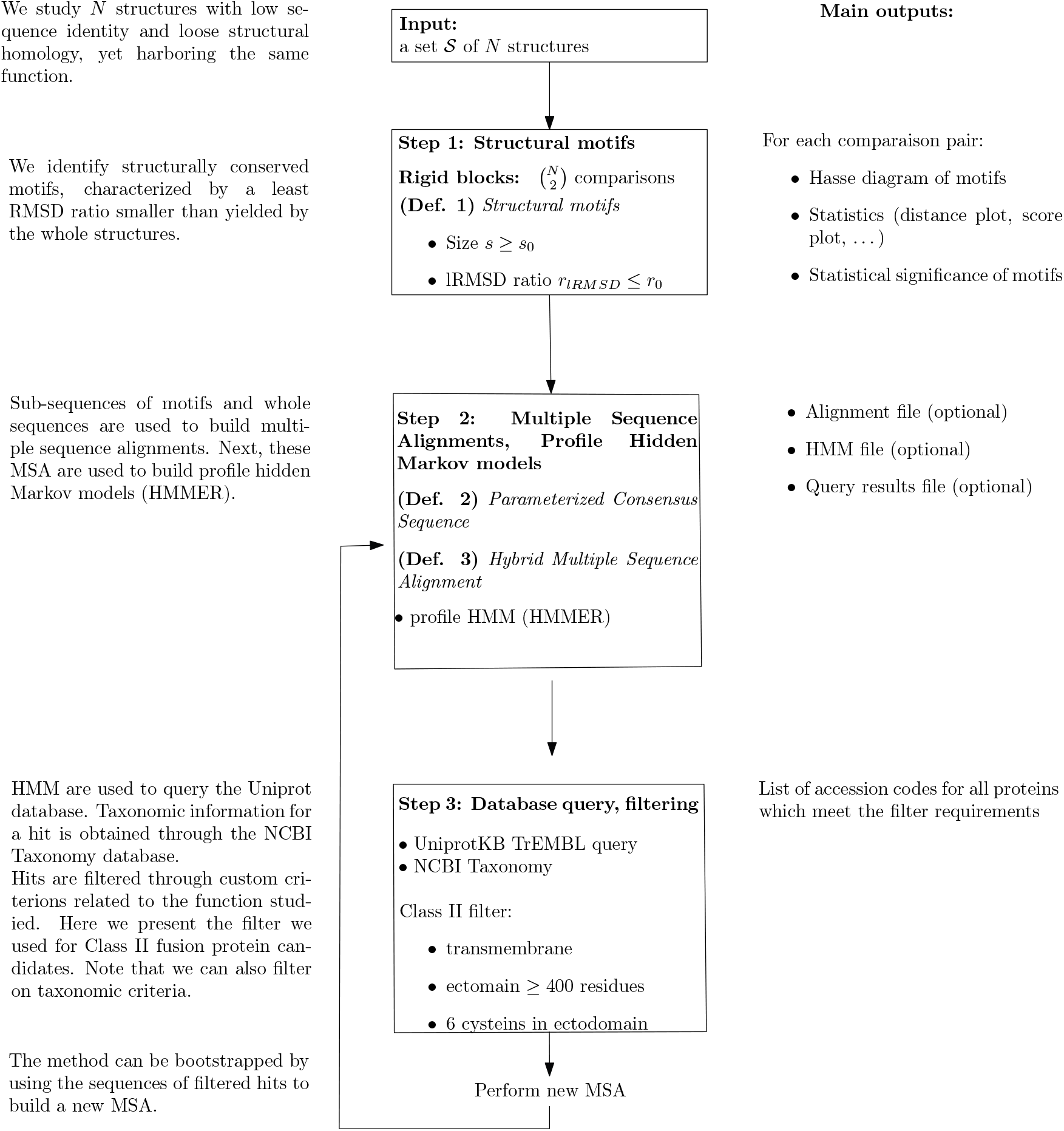
Sequence-structure based characterization of functionally related proteins: workflow.

- (Step 1) *Structural motifs* defined from pairwise structural alignments are collected. A motif harbors a lRMSD significantly smaller than that associated with a structural alignment between its two defining structures-the ratio between these two lRMSD is called the *lRMSD ratio*.
- (Step 2) Sub-sequences associated to motifs together whole protein sequences are used to build multiple sequence alignments (MSA). Combining sub-sequences and whole sequences is meant to bias the MSA towards the structurally conserved regions-deemed important, while yet retaining information on linkers connecting these regions. The MSA are then used to build profile HMM.
- (Step 3) HMM are used to query `UniProtKB`. The obtained hits are filtered, so as to retain the sequences with properties related to the function studied.

### 3.2 Step 1: Structural motifs

Given a set 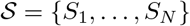 of *N* polypeptide chains, we define motifs on a pairwise basis.

#### Structural motifs for two structures

A large lRMSD between two structures possibly obliterates regions of smaller size and with a smaller lRMSD. To find such regions, given two structures, we define:

##### Definition. 1

*Consider two structures S_i_ and S_j_, and assume that a structural alignment between them has been computed. Given the two sets of a.a. identified by this alignment, i.e. M_i_ ⊂ S_i_ and M_j_ ⊂ S_j_, we define the least RMSD ratio as follows:*

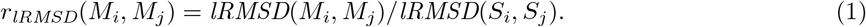

*The sets M_i_ and M_j_ are called structural motifs provided that* |*M_i_*| = |*M_j_*| ≥ *τ_MS_ and r_lRMSD_*(*M_i_*, *M_j_*) ≤ *τ*_/_, *for appropriate thresholds τ_MS_ and τ_/_*.

A black box returning such motifs is taken for granted. A generic framework to identify motifs was recently proposed in a companion paper [24]. The strategy hinges on two ingredients, namely an initial alignment providing a distance difference matrix (DDM), and a topological analysis of so-called filtrations coding conserved distances in the structures. We tested the four instantiations developed in [24], and obtained similar results (data not shown).

In the sequel, we report results obtained with two methods, namely `Align-Kpax-CD` (uses `Kpax` [25] as aligner; builds a filtration from conserved distances), and `Align-Kpax-SFD` (uses `Kpax` [25] as aligner; builds a filtration from a space filling diagram).

#### Thresholds used

In the sequel, we use *τ*_MS_ = 20 and vary *τ*_/_ in the range 0.5 … 0.8. Additionally, we define *nuggets*, motifs with the more stringent thresholds *τ*_MS_ = 20 and *τ*_/_ = 0.5.

##### Remark 1

*Note that for a given structure, a structural motif is not necessarily connected, neither on the structure, nor on the sequence. This stems from the fact that a motif is defined upon performing a structural alignment [24]*.

#### Structural motifs for N structures

We collect motifs for all pairs of comparisons. Sorting those with lRMSD ratio less than the threshold *τ*_/_ yields the following list:

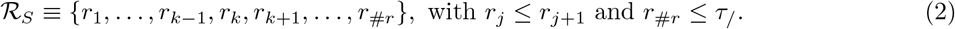

### 3.3 Step 2: From multiple sequence alignments to profile HMM

#### Parameterized consensus sequences

To exploit the sequence information of the motifs, we define a set of nested sub-sequences, parameterized by the lRMSD ratio found in the sorted list 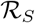 of Eq. (2):

#### Definition. 2

*(PCS) Consider a structure* 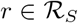 *together with a lRMSD ratio* 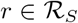. *The* parameterized consensus sequence 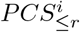 *associated with this structure is defined as the sequence of this structure, into which every amino-acid position not involved in any motif with lRMSD ratio less then r is replaced by a gap. The set of all* parameterized consensus sequences *is denoted PCS*.

Central in our method is the notion of PCS. On the one hand, stringent thresholds enforcing structural conservation are expected to yield motifs which may be too specific; on the other hand, too lenient thresholds may yield motifs which may lack specificity. Additionally, the structures present in 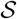 condition the motifs retrieved; in turn, the consensus sequences associated with these motifs are expected to be more or less specific of these sequences. This precisely motivates the definition of the various groups of interest (section 2).

#### Multiple sequence alignments and profile HMM

Using whole sequences and PCS, we define two sets of Multiple Sequence Alignment (MSA): MSA^Seq.^, a MSA involving the input protein sequences only; MSA^Hyb.^, a MSA involving the union of the protein sequences and the parameterized consensus sequences PCS. Practically, we use two multiple sequence aligners: ClustalΩ[26] and `MUSCLE` [27].

We exploit these MSA using profile hidden Markov models [5, 28] and the `HMMER` implementation [29, 30]. That is, to study the bias imposed by the addition of PCS to the full sequences, we define HMM^Seq.^ based upon MSA^Seq.^, and MSA^Hyb.^ based upon HMM^Hyb.^. These HMM models, whose complexity is measured by the number of match states, are used to perform database queries.

### 3.4 Step 3: Database queries and filtering

Database queries. A HMM is used to query `UniProtKB`. Such a query return *hits* in the form of `UniProtKB` accesion codes (unique identifiers tied to a protein sequence). The significance of hits being assessed with e-values (SI 7.4.4). For a given hit, we retrieve the taxonomic information from the `NCBI Taxonomy` database [31] (SI Sect. 7.4.4.) Note that we focus on species rather than protein sequences, since fragments, isoforms, or variants of a given protein typically correspond to separate entries.

#### Filtering hits using annotations: generic and fusion class II specific

The hits obtained can also be filtered using various criteria, which we illustrate for class II fusion proteins.

A first filter identifies transmembrane proteins. To find a transmembrane region in a hit from its FASTA sequence, we use Phobius [32]. The second filter aims at identifying class II candidates, based on three conditions: (i) at least one transmembrane region, (ii) an ectodomain (protein region extending into extracellular space) involving ≥ 400 residues, (iii) at least six cysteins in the ectodomain.

#### Cross-validation using HMM-HMM comparisons

Sequences obtained from `UniProtKB` can be rescored via HMM-HMM comparison (`HHpred`, [6]; see also SI Section 7.4). This strategy being of special interest to check the coherence with a HMM associated with a structure, we practically launch a query on the target database PDB_mmCIF70 (see https://toolkit.tuebingen.mpg.de/#/tools/hhpred).

##### Remark 2

*With respect to our overarching goal, which is to annotate sequences from* `UniProtKB`, *note that we use* `HMMER` *rather than* `HHpred`, *since since* `HHpred` *searches selected databases (PDB*, `Pfam`, *SMART, …) rather than* `UniProtKB`. *See* https://toolkit.tuebingen.mpg.de/#/tools/HHpred.

#### Bootstrap

The previous calculation may be integrated into a bootstrap strategy, with the sequences of the filtered hits incorporated so as to define new HMM (SI Fig. 6). We apply this strategy by running three bootstrap iterations on each previously defined groups with HMM^Hyb.^ and HMM^Seq.^. Note in passing that only sequences which yield an e-value ≤ 0.01 are considered for the bootstrap step.

#### Comparison of HMM^Hyb.^ and HMM^Seq.^: protocol

Summarizing the previous discussion, we compare HMM^Hyb.^ and HMM^Seq.^ as follows: (i) Fix a group of input sequences and structures, (ii) Fix a threshold *τ*_/_ to define structural motifs – Eq. (2)), (iii) Iteratively build the HMM models, perform the queries, count the species yielded, (iv) Bootstrap.

### 3.5 Software

The software implementing our methods is available in the Structural Bioinformatics Library [33] at http://sbl.inria.fr). The two main packages involved are StructuraLmotifs for the detection of motifs (https://sbl.inria.fr/doc/Structural_motifs-user-manual.html), and `FunChaT` for the functional characterization of proteins (https://sbl.inria.fr/doc/FunChaT-user-manual.html).

## 4 Results

### 4.1 Structurally conserved motifs

#### On the structural conservation of SSE

Class II proteins have a hierarchical structure, with SSE defining 23 structural units (SI Table 5), and a natural question is therefore to check whether the simplest structural elements, i.e. SSE, harbor any structural information. As established using hierarchical clustering methods, the short short answer is no (SI Sec. 7.3; SI Figs. DI: Fig. 7, DII: 8, DIII:9). This prompts for further analysis seeking structural conservation beyond SSE.

#### Motifs: structure and sequence conservation

To go beyond SSE, we used structural motifs yielded by the aforementioned algorihtm `Align-Kpax-CD` [24]. With thresholds τ_MS_= 20 and *τ*_/_ = 0.8, our method detects 188 structural motifs with sizes ranging from 20 to 116 (SI section 7.5) and lRMSD ranging from 0.63 to 10.73. Note that there can be redundancies (motifs that show little variation with respect to their constituting residues - typically less than 5). In using more stringent values (*τ*_MS_ = 20 a.a. and lRMSD ratio *τ*_/_ = 0.5), 118 structural motifs remain. Out of the 118, we handpicked 28 to minimize redundancies (SI Table 4 and SI Fig. 10). These nuggets are characterized by a size range 20… 67 residues, and lRMSD ratio range 0.09, …, 0.5 (SI Table 4).

From a functional standpoint, motifs may contribute indirectly (e.g. in defining the fold) or indirectly to the function of class II proteins. Various functional features of such proteins have indeed been characterized [16, 18, 19, 34]. Or critical importance is the hydrophobic fusion loop which is inserted into the target membrane, as well as the disulfide bonds stabilized the two loops emanating from the central domain I.

Closer inspection of our motifs reveals two important features (Fig. 3). On the one hand, several motifs sandwich (half)disulfide bonds. On the other hand, a motif typically spans several SSE elements (Fig. 3(Bottom)). This shows the difficulty of identifying such structurally conserved regions, as selecting the combination of SSEs together with their sub-components faces a combinatorial explosion-which we handle using geometric and topological techniques [24].

**Figure 3:**
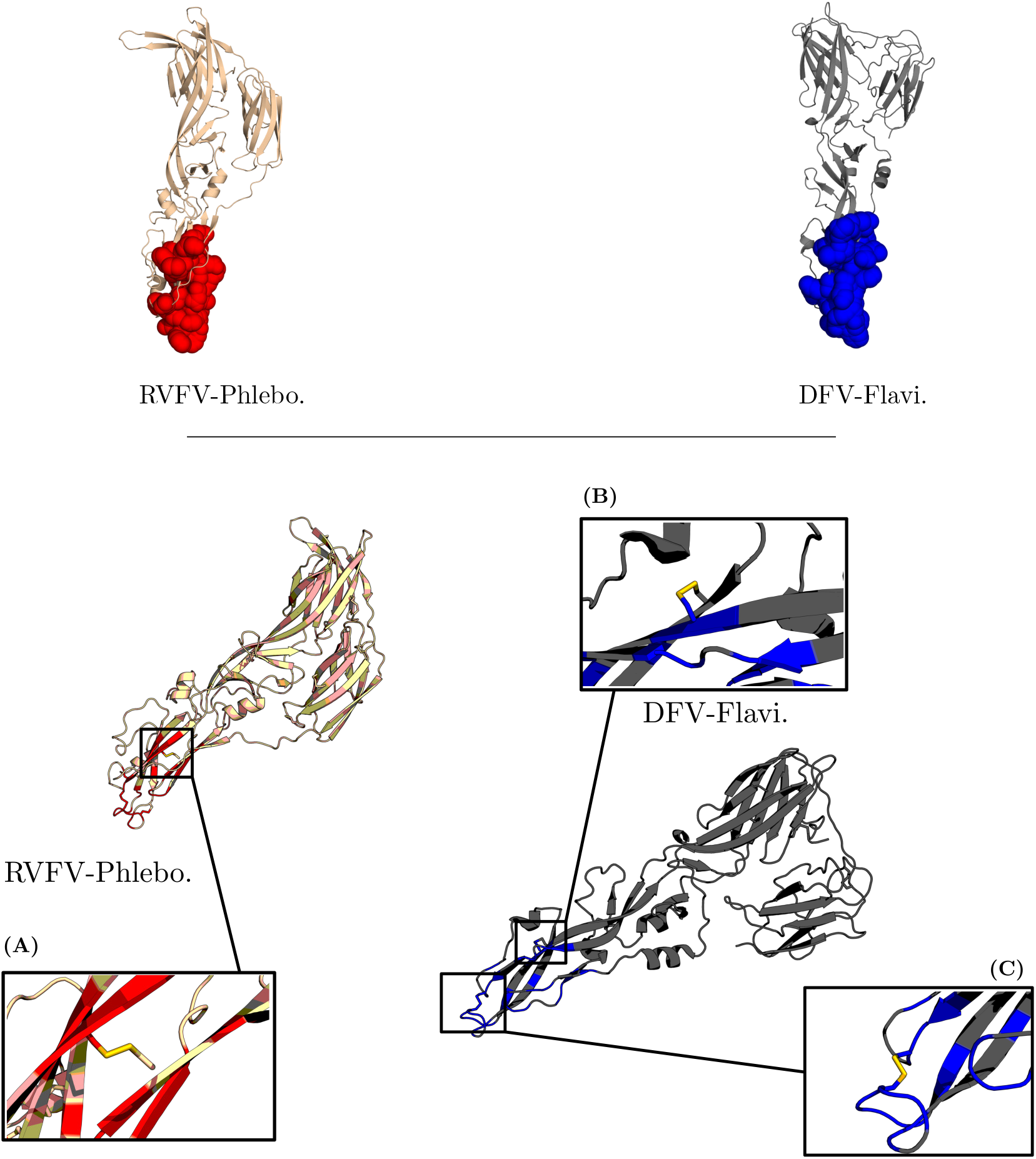
Comparing RVFV-Phlebovirus to DFV-Flavivirus: structural motifs. (**Top**) Motif represented with a solvent accessible model; the motif is localized on the tip of DII, which contains several disulfide bonds. (**Bottom**) Zoom on the motif, displaying the motif itself (red and blue amino-acids, respectively, on the two molecules), and the disulfide bonds within the motif.

### 4.2 Performances of hybrid HMMs for sequence retrieval

As a first assessment, since our hybrid HMM exploit sequence and structure information, we compare them to pure sequence based HMM to identify relevant sequences within `UniProtKB`. By varying the learning set and the threshold used to define structural motifs, we compare the species containing class II candidates retrieved over iterations. For these experiments, MUSCLE [27] is used to build the multiple sequence alignments.

More specifically, we compare the number of species identified both by HMM^Hyb.^ and HMM^Seq.^ (dark blue), those exclusively identified by HMM^Hyb.^ (orange), and those identified by HMM^Seq.^ (light blue) (Fig. 4, SI Sect. 7.4). Additionally, we investigate the complexity of the models (Table 2). Alongside the number of species, the hatched bars correspond to the number of emit states in a given HMM.

**Figure 4:**
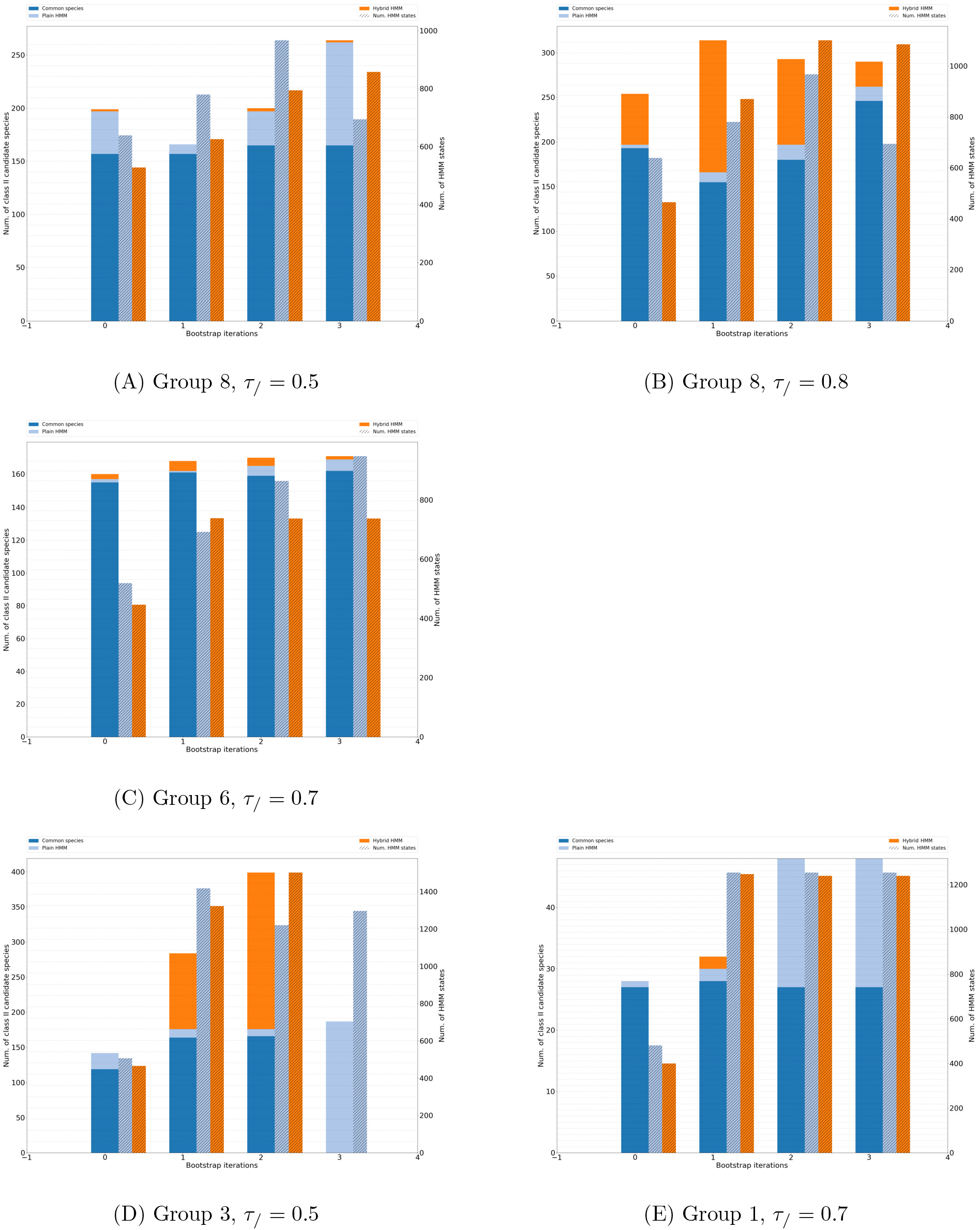
Various scenarios of domination when bootstrapping HMMs. For each iteration (0 to 3, x-axis), the 3 bars read as follows: first bar: species found by HMM^Seq.^ and HMM^Hyb.^ (solid blue), species found by HMM^Seq.^ only (light blue), species found by HMM^Hyb.^ only (orange); second bar: hatched light blue: number of emit states of HMM^Seq.^; third bar (hatched orange): number of emit states of HMM^Hyb.^. **(A)** HMM^Seq.^ consistently dominates HMM^Hyb.^. **(B)** HMM^Hyb.^ consistently dominates HMM^Seq.^. **(C)** Both types of HMM yield comparable number of specific species. **(D)** At each bootstrap iteration, HMM^Hyb.^ shows a large increase in number of species until the model becomes too complex and the HMM implementation used fails to manage it. **(E)** HMM^Seq.^ displays a peak number of species at the second bootstrap iteration.

General comments are in order:

- In general, both models provide specific information i.e. species that are not found by the other model (Fig. 4; all figures, SI Sect. 7.6).
- Overall, both methods report comparable number of species (Table 1), with HMM^Hyb.^ slightly more stable than HMM^Seq.^ (std. dev. of 271 versus 290). This overall observation hides cases where each model alternatively dominates the other in a stable fashion (Fig. 4(A,B) and cases where both models fare equivalently (Fig. 4(C)).
- Bootstrap iterations are either characterized by a sharp rise of the number of species reported (Fig. 4(D,E)), or by a rather stable behavior (Fig. 4(A,B,C)).
- The performances of HMM^Hyb.^ are conditioned to the set of motifs used, which is itself parameterized by the ratio *τ*_/_ – Eq. (2). Stringent threshold indeed result in a smaller bias imposed by structural motifs. At threshold (*τ*_/_ = 0.5), it appears that only group 3 and 7 enjoy unique species identified (SI Fig. 12). Beyond that threshold, no clear rule emerges to select *τ* (SI Figs. 13, 14, 15).
- In terms of model complexity, both HMM^Hyb.^ and HMM^Seq.^ fare equally. However, HMM^Hyb.^ is, on average, slightly smaller at the initial step, and grows slightly larger in the final stages (Table 2). From the first bootstrap step and on-wards, the model sizes nearly double.

**Table 1:** Retrieved species: statistics over the four runs and all *τ*_/_ thresholds determining motifs. Statistics are reported over four runs i.e. the initial run + three bootstrap iterations; three values of parameter *τ*_/_, which determines structural motifs, were used: *τ*_/_ = (0.5,0.6, 0.7,0.8). Species variations refers to the variation in-between two consecutive runs.

| Model | # species |  | # species variation |  |
| --- | --- | --- | --- | --- |
| | Mean | $\sigma$ | Mean | $\sigma$ |
| HMM<sup>Hyb.</sup> | 284 | 271 | 103 | 188 |
| HMM<sup>Seq.</sup> | 283 | 290 | 118 | 222 |

**Table 2:** HMM complexity: size of the model i.e. number of emit states on each bootstrap iteration. Initially, HMM^Hyb.^ is slightly smaller and more stable than HMM^Seq.^. In later stages, the opposite behavior is observed.

| Model | HMM num. states |  |  |  |  |  |  |  |
| --- | --- | --- | --- | --- | --- | --- | --- | --- |
|  | Initial model |  | First bootstrap |  | Second bootstrap |  | Third bootstrap |  |
| | Mean | $\sigma$ | Mean | $\sigma$ | Mean | $\sigma$ | Mean | $\sigma$ |
| HMM<sup>Hyb.</sup> | 479 | 59 | 1319 | 886 | 1383 | 754 | 1275 | 530 |
| HMM<sup>Seq.</sup> | 558 | 100 | 1351 | 876 | 1356 | 757 | 1163 | 328 |

To further these insights, we inspects characteristic scenarios (Fig. 4).

- Even though both types of models are relatively stable, the complexity of the model may *explode* at a given iteration, inducing a net increase in the number of species found and also potentially noise. The implementation of HMM may be unable to handle a large number of emit states (Fig. 4(A), HMM^Hyb.^, 3rd iteration).
- Adding or removing sequences from a model can have any type of effect. Addition of new sequences to expand the MSA and the associated HMM is an expected behavior. But the opposite is also observed. Consider the case where the initial calculation is such that HMM^Hyb.^ does not retrieve any specific sequence (Fig. 4(D), leftmost column). Therefore, the sequences used to build the MSA for HMM^Hyb.^ are a strict subset of the sequences used to build the MSA for HMM^Seq.^. Yet, at the first bootstrap iteration, HMM^Hyb.^ retrieves more species than HMM^Seq.^ (Fig. 4(D), first column).

#### Remark 3

*Upon investigation some hits do not meet the requirements of our filter because the sequence is partial–regions containing the cysteins and/or the trans-membrane region may be missing. This reflects poorly on group 1, which displays very little species. In reality, both HMMs are able to recover most of the species of its corresponding taxonomic group*.

### 4.3 Performance of hybrid HMMs to retrieve homologs of HAP2-GCS1

We noted in Introduction that fusogens in viruses and eukaryotes evolve at different speeds, due in particular to the selection pressure imposed to the former by immune systems. As a second assessment of our HMMs, we therefore narrow down our focus on eukaryotes, and study the ability of our hybrid HMM to identify homologs of HAP2-GCS1.

#### Species

With a focus on eukaryotes irrespective of the differences between EFF-1 and HAP2-C, we further investigate performances of both HMM models using Group 4 at threshold *τ*_/_ = 0.7. This group was chosen because the HAP2-GSC1 family is currently of high interest; a general aim is to find members of this family among larger organisms, such as vertebrates. The *τ*_/_ = 0.7 threshold seems to yield the best results for HMM^Hyb.^. We wish to estimate how well the models characterize the HAP2-GSC1 protein family. To do this, we exploit `UniProtKB` sequence annotations (http://www.uniprot.org/help/sequence_annotation) to find hits which have been labeled as having a HAP2-GSC1 domain. At the initial step, HMM^Hyb.^ recovers 254 hits spread across 149 species; HMM^Seq.^ finds 244 hits (2 of which are exclusive) across 145 species. At the third and final bootstrap iteration, HMM^Hyb.^ 228 hits across 126 species; HMM^Seq.^ finds 298 hits across 167 species. In this case, HMM^Hyb.^ finds 12 exlusive hits across 10 species. The following remarks are in order:

- A very small training set is enough to recover many homologs (for both models).
- Initially HMM^Hyb.^ performs better at finding HAP2-GSC1 family members.
- After the bootstrap iteration, HMM^Seq.^ performs better although HMM^Hyb.^ still has exclusive species.

Note that this tally is not exhaustive as some proteins that are known HAP2-GSC1family members are not annotated. For example, the Tetrabaena socialis HAP2 protein (`UniProtKB` accession code A0A2J8AI85), does not have the HAP2-GSC1 domain annotation. Even though it is a know HAP2 protein, it will not show up in the HAP2-GSC1 domain count. Additionally, members of HAP2-GSC1 in larger organisms (such as vertebrates) are theorized to be distant homologs. A model that is too specific could be a hindrance to finding such a protein so that loosing some hits in the further steps should not be necessarily seen as an obstacle torward that goal.

#### Homologs of HAP2-GCS1

To further constrain the models, we restrict the learning set to the domains II of three HAP2-GSC1 structures known at the time of this study. The three structures accession numbers are 5MF1 (HAP2e) [22], 5OW3 (AtHAP2) and 5OW4 (TcHAP2) [23]. Using motifs at threshold *τ*_/_ = 0.8, we build a new HMM^Hyb.^ (SI Fig. 11 for its sequence logo.) When results are displayed, the particular method used is indicated (by method we refer to `Align-Kpax-CD`, `Align-Kpax-SFD`, see Section 3.2, and the multiple sequence alignment used).

To assess the ability of our method to find remote homologs, we check for the recovery of the most distant known homolog to our training set, the HAP2-GCS1 in the drosophila fly. It was indeed shown [35] that D. melanogaster gene CG34027, an ortholog of HAP2-GCS1, codes for a protein involved in plasma membrane fusion. Different combinations of filtration methods as well as sequence aligners yield different results (Tab. 3). From the first iteration, we recover a number of known HAP2 sequences, notably in arthropodes (34 species, data not shown). From the first bootstrap iteration, we recover 17 plausible HAP2-GCS1 candidate sequences in 10 different Drosophila species. We cross validate these results using `HHpred` (Sect. 3.4 on HMM-HMM comparisons). Of particular interest are comparisons against the known gamete fusion protein in chlamydomonas reinhardtii (HAP2 structure, pdbid 5MF1). Eleven of the sequences returned a very low e-value leading to the conclusion that they are most probably HAP2 proteins (Tab. 3). One of these corresponds to the aforementioned gene CG34027 (`UniProtKB` identifier: Q2PDQ0).

**Table 3:** Searching for remote HAP2-GSC1 homologs: hits in the drosophilia fly. Cross-validation of the sequences yielded by our method –see Section 3.4. Reported are the top hits obtained with `HHpred` (3rd column), with small e-values indicating a likely HAP2 protein (4th column). The pdbids associated with the hits are also given; those corresponding to known HAP2 structures are marked in bold.

| Species | UniProtKB accession | HHpred top hit (PDB ID) | e-value |
| --- | --- | --- | --- |
| **Align-Kpax-CD, ClustalΩ** |  |  |  |
| *Drosophila ananassae* | B3M263 | 5MF1 | 3e-51 |
| *Drosophila mojavensis* | A0A0Q9X6U7 | 5MF1 | 8.7e-55 |
| *Drosophila virilis* | A0A0Q9WDI4 | 5MF1 | 2.1e-53 |
| **Align-Kpax-CD, Muscle** |  |  |  |
| *Drosophila mojavensis* | A0A0Q9X6U7 | 5MF1 | 8.7e-55 |
| *Drosophila virilis* | A0A0Q9WDI4 | 5MF1 | 2.1e-53 |
| **Align-Kpax-SFD, ClustalΩ** |  |  |  |
| *Drosophila ananassae* | B3M263 | 5MF1 | 3e-51 |
| *Drosophila mojavensis* | A0A0Q9X6U7 | 5MF1 | 8.7e-55 |
| *Drosophila virilis* | A0A0Q9WDI4 | 5MF1 | 2.1e-53 |
| **Align-Kpax-SFD, Muscle** |  |  |  |
| *Drosophila erecta* | B3P6S0 | 5MF1 | 2.7e-39 |
| *Drosophila sechellia* | B4IJG1 | 5MF1 | 2.7e-39 |
| *Drosophila ananassae* | B3M263 | 5MF1 | 3e-51 |
| *Drosophila melanogaster* | Q2PDQ0 | 5MF1 | 6.5e-31 |
| *Drosophila mojavensis* | A0A0Q9X6U7 | 5MF1 | 8.7e-55 |
| *Drosophila mojavensis* | A0A0Q9WXE1 | 5MF1 | 1.5e-61 |
| *Drosophila mojavensis* | A0A0Q9WWX9 | 5MF1 | 1.4e-67 |
| *Drosophila yakuba* | B4PUD6 | 5MF1 | 1.3e-32 |
| *Drosophila mojavensis* | A0A0Q9WX02 | 5MF1 | 1.6e-45 |
| *Drosophila ficusphila* | A0A1W4VL29 | 5MF1 | 1.8e-52 |
| *Drosophila virilis* | A0A0Q9WDI4 | 5MF1 | 2.1e-53 |
| *Drosophila virilis* | A0A0Q9WEW4 | 5IJ0 | 3.7 |
| *Drosophila ananassae* | A0A0P8YDK7 | 3S84 | 10 |
| *Drosophila mojavensis* | A0A0Q9X9J0 | 3S84 | 9.3 |
| *Drosophila mojavensis* | B4KU79 | 3S84 | 8.4 |
| *Drosophila simulans* | A0A0J9RKJ6 | 3S84 | 8.5 |
| *Drosophila pseudoobscura pseudoobscura* | Q28WV3 | 3S84 | 9.5 |

Importantly, none of these eleven sequences mentions the Pfam domain PF10699 (https://pfam.xfam.org/family/PF10699). This fact illustrates the ability of our structural models to capture structural information which is more localized than whole Pfam domains.

## 5 Discussion and outlook

Remote homologous proteins are proteins sharing low sequence identity, yet having similar structures and functions. As the development of sequencing techniques yields a rapid increase in the number of known sequences, while the number of solved structures grows more slowly, the ability to detect remote homology is a central problem in structural bioinformatics. Our work contributes a novel method for this problem, combining sequence-structure information. As an application, we focus on the problem of identifying within `UniProtKB` sequences which might be class II fusion proteins with high probability. Using a diverse learning set involving structures from viruses and eukaryotes, we show that our hybrid HMM models retrieve proteins from species which are not identified from pure sequence based HMM. Moreover, given the relative diversity of structures for class II fusion proteins, we also show that using a narrower learning set involving eukaryotic structures only, our method identifies remote homologs in D. melanogaster, which are not retrieved using the relevant PFAM domain.

To discuss the merits of our method, it is informative to consider in turn the three types of information a remote homology detection method may enjoy: sequences, structures, and dynamic information.

When sequences and/or structures of proteins with a common function are known, the classical route consists in modeling these sources of information with multiple sequence alignments and/or profile HMM and/or ad hoc feature spaces. Along this line, our contribution is to show that biasing profile HMM with structural features detected amongst a learning set (a handfull of structures) indeed helps to identify novel sequences compatible with the function targeted. Further cross-validation of the sequences retrieved via HMM-HMM comparison against witness HMM models with known structures provides a high level of confidence. However, this strategy is only partially satisfactory, since sequences that do not yield a low e-value via HMM-HMM comparison may still be of high interest.

Our ability to retrieve remote homologs of HAP2 in drosophilia melanogaster stresses the relevance of the structural motifs underlying our hybrid HMM, which typically span portions of SSE elements, and whose detection is a non trivial endeavor. We also note that biasing profile HMM with structural information, albeit a straightforward idea, is a rather subtle strategy for two reasons. On the one hand, biased HMM models, when compared to pure sequence based models, do bring unique features-as evidenced by the fact that they are the only ones to identify selected sequences from `UniProtKB`; yet, both classes of models have unique traits since they do identify unique species. On the other hand, the learning set plays a crucial role in particular in terms of diversity, and qualifying the output as a function of this diversity remains an open problem.

So far, we have excluded from the design of remote homology detection methods information related to the dynamics, including structural (meta-stable states), thermodynamic (occupancy probabilities), and kinetic information (transition rates between states). In the presence of large amplitude conformational changes, this type of information is admittedly out of reach in most cases for experimental and simulation methods. However, amino-acids involved in key structural, thermodynamic or dynamic events would naturally help improving all types of remote homology detection methods (alignment methods, discriminative methods, ranking methods), and would also alleviate the aforementioned cross-validation step in the absence of obvious witnesses. More generally, such insights might dramatically reduce the region of sequence space in which distant homology is sought. In fact, in moving from sequence to dynamics across structures, one uses finer mechanism related information, which is however more challenging to get. Finding the optimal combination appears as a very promising research avenue, calling for deep insights on the connexions between sequence, structure, dynamics and function.

